# *Scaphoideus titanus* Ball feeding behaviour on three grapevine cultivars with different susceptibilities to Flavescence dorée

**DOI:** 10.1101/2021.11.25.470030

**Authors:** Matteo Ripamonti, Federico Maron, Daniele Cornara, Cristina Marzachì, Alberto Fereres, Domenico Bosco

**Author notes:** Environmental Research and Innovation Department (ERIN), Luxembourg Institute of Science and Technology (LIST), Esch-sur-Alzette, Luxembourg. Vit.En. Centro di Saggio, Calosso (AT), Italy. University of Berkeley, Department of Environmental Science, Policy, and Management, Berkeley, CA, USA.

## Abstract

*Scaphoideus titanus* (Ball) is a grapevine-feeder leafhopper, and the most important vector of Flavescence dorée of grapevine (FD), a disease associated with phytoplasmas belonging to ribosomal subgroups 16Sr-V–C and –D. FD is a major constraint to viticulture in several European countries and, so far, its control has relied on roguing of infected plants and insecticide applications against the vector. Detailed knowledge on different levels of the multifaceted phytoplasma-plant-vector relationship is required to envisage and explore more sustainable ways to control the disease spread. In the present work, *S. titanus* feeding behaviour was described on three grapevine cultivars: Barbera (susceptible to FD), Brachetto, and Moscato (tolerant to FD) using the Electrical Penetration Graph (EPG) technique. Interestingly, no differences were highlighted in the non-phloem feeding phases, thus suggesting that the tested cultivars have no major differences in the biochemical composition or structure of the leaf cuticle, epidermis or mesophyll, that can affect the first feeding activities. On the contrary, the results showed significant differences in leafhopper feeding behaviour in terms of the duration of the phloem feeding phase, longer on Barbera and shorter on Brachetto and Moscato, and of the frequency of interruption-salivation events inside the phloem, higher on Brachetto and Moscato. These findings indicate a possible preference for the Barbera cultivar, a better host for the leafhopper. *Scaphoideus titanus* feeding behaviour on Barbera correlates with an enhanced FDp transmission efficiency, thus explaining, at least in part, the higher susceptibility of this cultivar to FD. The mechanisms for the possible non-preference for Brachetto and Moscato are discussed, and an antixenosis is hypothesized. We propose that breeding for resistance against FD should take into account both plant traits associated with the response to the phytoplasmas and to the vector.

## 1. Introduction

The leafhopper *Scaphoideus titanus* (Ball, 1932. Hemiptera: Cicadellidae: Deltocephalinae) is the main vector of phytoplasmas associated with the Flavescence dorée of grapevine (FD), a disease spread in most European viticultural countries (European Food Safety Authority, 2020) that causes severe reduction of yield and quality of grapes, requires roguing of infected plants and leads to uneven-aged vineyards (Morone et al., 2007). FD is associated with phytoplasma agents belonging to the 16SrV group, subgroups –C and –D (Davis and Dally, 2001; Lee et al., 2004; Martini et al., 2002), and it causes severe losses to European viticulture (EFSA, 2016). Although different insect species are competent for the transmission of FD phytoplasmas (FDp), *S. titanus* is by far the most important vector, being strictly associated with *Vitis* plants and thus sustaining both primary (from wild grapevines outside the vineyards to cultivated vines) and secondary (from vine to vine within the vineyard) disease spread (Maggi et al., 2017; Ripamonti et al., 2020). Control of FD relies on prophylactic measures, such as the use of healthy propagation material, as well as on compulsory measures in infected vineyards, namely roguing of infected plants, and insecticide treatments against the vector (Bosco and Mori, 2013). However, the large-scale application of insecticides is a concern to human health and environment, priming cascade ecosystem effects (Desneux et al., 2007) with a strong negative impact on pollinators (Tosi et al., 2018). For this reason, recent studies focused on identifying sources of resistance to FDp phytoplasmas within the grapevine germoplasm (Eveillard et al., 2016; Ripamonti et al., 2021), that would represent the best strategy to minimise damage and limit FD spread and insecticide applications. Grapevine tolerance to FDp may be due to a direct response of the plant against the pathogen or mediated by some resistance against the vector, or by a combination of the two. Resistance against insects occurs when plant structural or chemical traits deter herbivore feeding and thus minimize the amount of herbivore damage experienced by the plant, while tolerance occurs when plant traits reduce the negative effects of herbivore damage on crop yield (Mitchell et al., 2016). As an example, it was demonstrated that resistant tea cultivars sustained lower phloem ingestions for *Empoasca vitis* (Miao et al., 2014). Moscato and Brachetto are grapevine cultivars tolerant to FD, as demonstrated by Ripamonti et al. (2021) using transmission experiments with *S. titanus* under controlled conditions (Ripamonti et al., 2021). The reduced *S. titanus* survival on Moscato observed by the above mentioned authors, suggest that vector-host interaction could be the pivotal factor underlying Moscato tolerance to FD. *S. titanus* is monophagous on *Vitis* species, mainly *Vitis vinifera* and naturalized rootstocks of *V. riparia* in Europe, while in North America, *V. labrusca* and *V. riparia* are reported as the preferred host plants (Chuche and Thiéry, 2014). Although the species is regarded as monophagous, it shows a good level of plasticity and can feed on plant species of different families, e.g. Vitaceae, Fabaceae, Ranunculaceae (Caudwell et al., 1970; Trivellone et al., 2013). Plant resistance against sap-sucking insects can be conveniently investigated by Electrical Penetration Graph (EPG), that describes the feeding behavior of a sucking insect on a given plant genotype, by identifying possible altered nutrition on non-suitable genotypes (Backus et al., 2020; Lucini et al., 2021).

Here we expand the first findings on *S. titanus* behaviour on grapevines, by analyzing the vector probing behavior on three cultivars with a different degree of susceptibility to FD: one susceptible, Barbera, and two tolerant, Moscato and Brachetto, through the Electrical Penetration Graph (EPG) (Backus and Bennett, 2009; McLean and Kinsey, 1964; Tjallingii, 1978).

EPG is a powerful tool to describe pierce-sucking insects’ probing behaviour, previously applied to describe *S. titanus* feeding behaviour on Cabernet-Sauvignon cuttings (Chuche et al., 2017a, 2017b). EPG studies on different plant cultivars/genotypes provide precious information for the epidemiology of vector-borne plant pathogens, also permitting the identification of traits making a *Vitis* genotype unsuitable for the vector. A number of EPG studies aimed at identifying plant resistance to insect vectors have been performed on planthoppers (Kimmins, 1989), whiteflies (Jiang et al., 2001; Rodríguez-López et al., 2011) and aphids (Caillaud et al., 1995a, 1995b; Garzo et al., 2018; Sauge et al., 1998). Among these latter, EPG was applied to identify resistance factors involved in virus transmission inhibition (Chen et al., 1997) as well as the presence of antixenosis (Kordan et al., 2019). Besides those on *S. titanus* (Chuche et al., 2017a, 2017b), few EPG studies have been conducted on Deltocephalinae leafhoppers (Carpane et al., 2011; Kawabe and McLean, 1980; Lett et al., 2001; Stafford and Walker, 2009; Trębicki et al., 2012), and very few of these are focused on phytoplasma vectors.

The aim of this study was to compare *S. titanus* feeding behaviour on three different grapevine cultivars with different susceptibilities to FD, in order to better characterize the mechanisms underlying varietal tolerance/susceptibility to this phytoplasma disease.

## 2. Material & Methods

### 2.1. *S. titanus* collection and rearing

To establish a *S. titanus* laboratory colony, in January/February 2019 two-year-old grapevine canes with eggs were collected in vineyards of the Piemonte Region. The selected sites were known to host a high population of the leafhopper in the previous summer, as estimated by yellow sticky traps captures of adults. The collected canes were stored in a cold room at 6±1°C, covered with plastic film to avoid egg desiccation, until use. When needed, grapevine canes were transferred into an insect-proof greenhouse at 24 ± 2°C and maintained damp by daily water spraying. After four weeks, canes were isolated in a cage together with a three-week-old broadbean plant as a food source for the nymphs. After egg hatching, the broadbean plant was replaced every three weeks. Nymphs were reared under controlled conditions inside a greenhouse chamber, at 24 ± 2°C, with no humidity and photoperiod control, from the beginning of April to the end of September 2019. As FDp is not transovarically transmitted, and all the plants used for the rearing and the experiments were phytoplasma-free, all *S. titanus* used in the experiments were phytoplasma-free. For the EPG experiments, adults from 7 to 21 days after emergence were used (modified from Chuche et al., 2017a), since in this time frame they were sexually mature, highly active and not subjected to high mortality (Bocca et al., 2020; Mazzoni et al., 2009).

### 2.2. Plant rearing

The test plants were obtained from phytoplasma-free *V. vinifera* cuttings of three different cultivars, Barbera N. - Clone I-AT 84, Brachetto N. - Clone I-CVT 20 and Moscato Bianco B. - Clone I-CVT 190 as described in Ripamonti et al. (2021). Grapevine cuttings were grown in a greenhouse at 24 ± 2°C, with no humidity and photoperiod control, inside 0.9 L pots (2:2:1 topsoil, clay, perlite), and watered once a week. Cuttings were three to five months old, pruned in order to keep the height within 80 cm. Broadbean plants used for *S. titanus* rearing were seedlings maintained in a growth chamber (24 ± 2°C, with no humidity and photoperiod control) in 2.4 L topsoil, five per pot, and watered twice a week.

### 2.3. EPG setup and data analysis

Selected adults were collected and anesthetised with carbon dioxide for 5 seconds in a glass tube, then immobilised at the edge of a cut pipette tip connected to a vacuum pump under a stereomicroscope. A small drop of water-based silver glue (EPG Systems, Wageningen, The Netherlands) was placed on the pronotum of the insect, then a gold wire of 18 μm (previously attached with solvent-based silver glue (Ted Pella Inc., USA) to a 3 cm copper wire in turn attached to a brass nail with melted stain) was positioned on the dried drop, and covered with another small drop of silver glue. Before the EPG assay, insects were starved for a 30-minute period, during which they were attached to the electrode and hanged, inserting the nail in a polystyrene base.

The substrate voltage probe was inserted in well damped soil of a potted grapevine cutting, and *S. titanus*, attached to the assembled electrode, was connected to a probe and positioned onto the abaxial surface of a leaf. The feeding behaviour was then monitored for 8 hours with a Giga-8dd DC-EPG amplifier (EPG Systems, Wageningen, The Netherlands), inside a Faraday cage to isolate the system from external electrical noise. Input resistance used was 1 giga Ohm, output set at 75x gain and plant voltage adjusted so that the EPG signal fitted into +5V and -5V. All recordings were done between June and August 2019, and started between 11:00 and 11:30 a.m. every day.

A total of 153 recordings were done, each day a total of 6 recording were run. Each single recording was represented by a different plant-insect combination, one male or one female on one grapevine plant. Potted plants of the three cultivars were randomly arranged in the Faraday cage for every recording and discarded after use. In case of falling from the leaf, the insect was repositioned. At the end of the recording, dead insects were noted and excluded from further analyses.

### 2.4. EPG acquisition and marking of EPG files

Recordings were acquired and marked using Stylet+ software (v01.30, Electrical Penetration Graph Data Acquisition and Analysis, EPG Systems, Wageningen, The Netherlands). Waveform marking was conducted accordingly to Chuche et al. (2017a) and Stafford & Walker (2009), focusing on the following behavioral phases: np (non-probing activity), pathway-phase (phase “C”), active ingestion (phase “G”) of mesophyll (<60 seconds) or xylem sap (>100 seconds) (see Stafford & Walker, 2009), passive ingestion of phloem sap (phase E), interruptions during ingestion (phase N of Chuche et al., 2017a). For more details, see Supplementary File S1.

Once marked, all the recordings were singly selected for the successive analysis. In particular, recordings with electrical noises, bad electric connections, or when insects fell from the plant for more than 20% of the recording time, were discarded from further analysis.

### 2.5. Statistical analysis

All the statistical analyses were conducted on R software v4.0.3 (R Core Team., 2020). Selected recordings were analysed through a package of the software R ad-hoc produced for the analysis on EPG recordings, called Rwaves (Chiapello et al., https://github.com/mchiapello/Rwaves; manuscript in preparation)). Rwaves conducts summary statistics on the input recordings on a set of variables of EPG analysis (Table 1), producing a table including the values of all the variables for all the input recordings. The resulting table was composed as follows: every row corresponded to a single recording (represented by the unique combination of one leafhopper and one grapevine plant), while every column represented a single EPG variable. Once obtained, the table was subjected to modifications to enhance readability (packages dplyr, tidyr, stringr: (Wickham, 2019, 2020; Wickham et al., 2020), and descriptive statistics were run (Tables 3, 4, 5, 6 and Supplementary File S2). Univariate analyses were conducted starting from Generalised Linear Model (GLM) of different families specific for the nature of the variable: quasi-Poisson or negative-binomial for counts, Gamma or inverse-Gaussian for positive continuous variables, beta-regression for proportions (packages stats, betareg, MASS: Cribari-Neto and Zeileis, 2010; Venables and Ripley, 2002). Goodness-of-fit for every model was evaluated plotting half-normal plots with simulated envelope against deviance residuals, with 95% confidence level (hnp package: Moral et al., 2017). Homoscedasticity for every model was evaluated through Levene’s test (car package: Fox and Weisberg, 2019). In case of rejection of the null hypothesis, heteroscedasticity-consistent standard errors (sandwich package: Zeileis et al., 2020) were calculated and considered for pairwise-comparisons. Comparisons among groups were conducted with least-square means method and Tukey method for p-value adjustment, at 0.05 significance level and 95% confidence intervals (packages emmeans and multcomp: Hothorn, Bretz, & Westfall, 2008; Lenth, 2020). Cultivar, Sex, and their reciprocal interaction were selected as explanatory variables. If no significant effects were found for Sex and Cultivar × Sex, the GLM was run with Cultivar as the only explanatory variable. GLMs summaries were reported in Supplementary File S3 using package jtools (Long, 2020). Packages ggplot2 (Wickham, 2016) and ggpubr (Kassambara, 2020) were used to produce Figure 1, and Supplementary File S4 and S5.

**Table 1.**
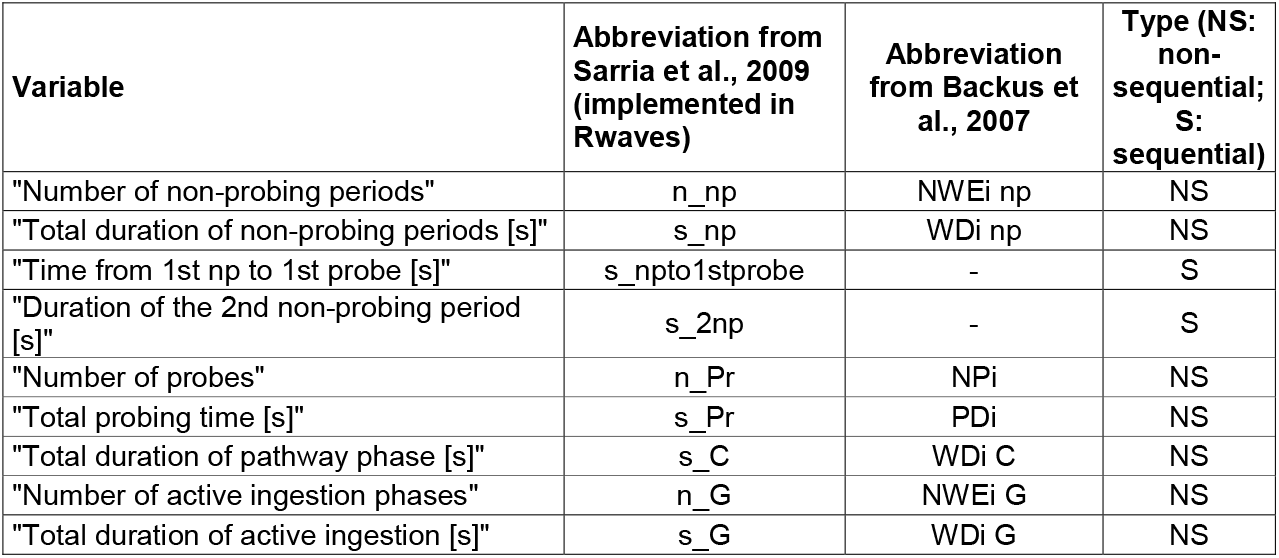
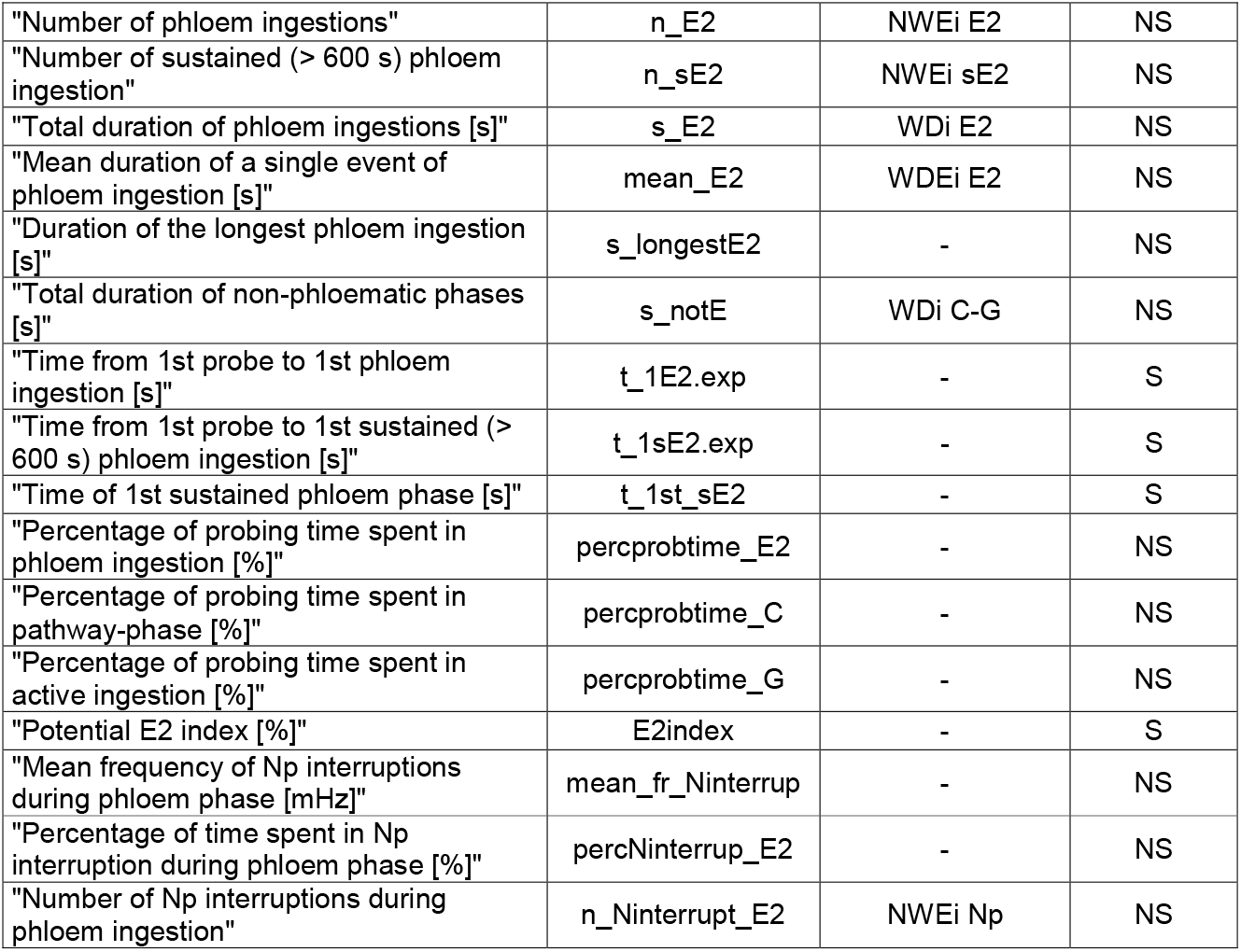
EPG variable selected for the study.

**Figure 1.**
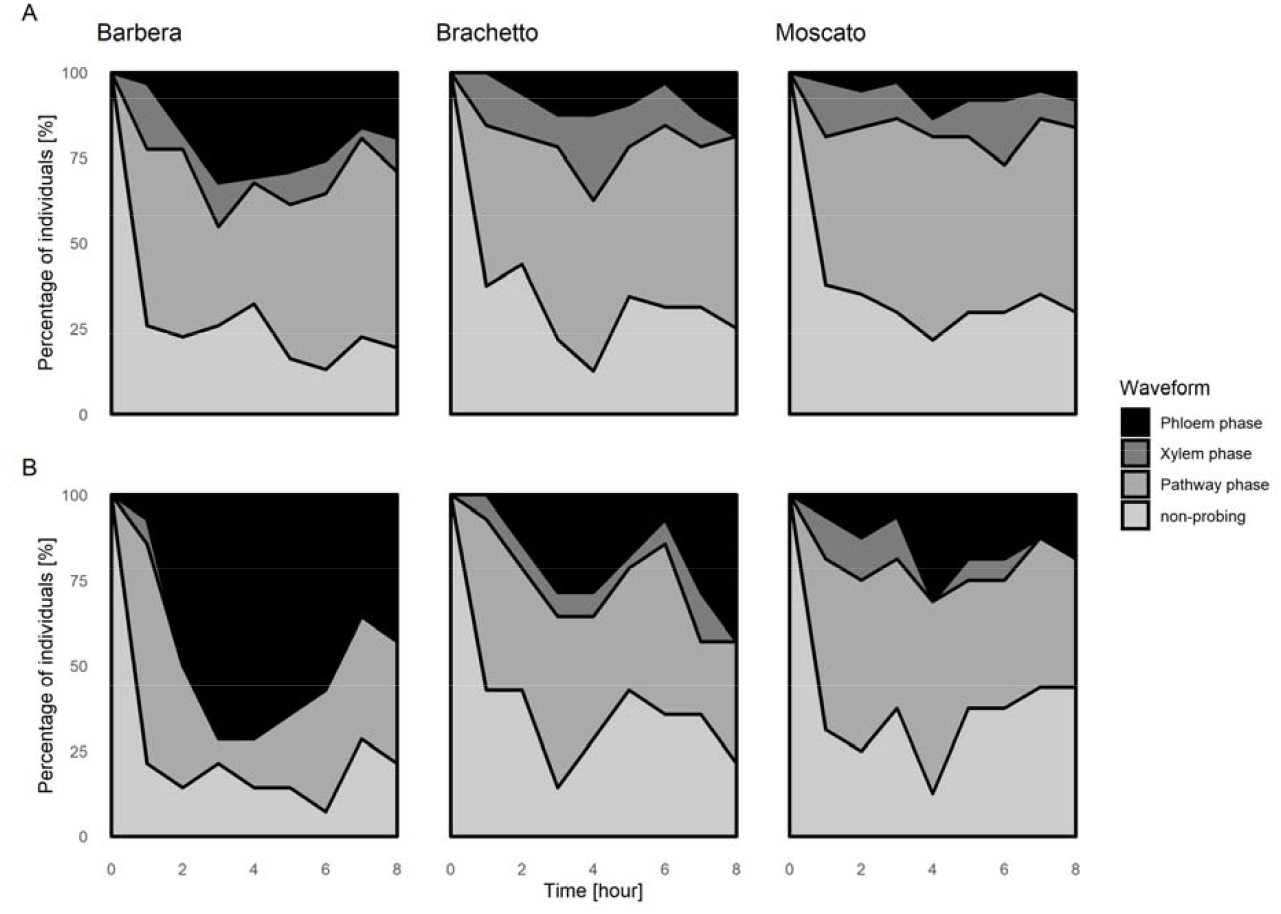
Temporal progress of *S. titanus* behavioral phases on three grapevine cultivars during the 8-h EPG recording. Probing behaviours were represented as percentages of leafhoppers in a given phase (non-probing, pathway phase, xylem phase, phloem phase) at 1 h intervals, starting from hour 0 (start of the recording) to hour 8 (end of the recording). a) Graphs produced considering all recordings; b) graphs produced considering only recordings where a phloem phase was present. The total number of recordings used to produce Figure 1 are reported in the third column of Supplementary File S2 (all recordings) and Table 3 (phloem recordings).

A multivariate Canonical Correspondence Analysis (CCA, Legendre & Legendre, 2012) was conducted through the vegan (Oksanen et al., 2019) and ggordiplots packages (Quensen, 2018), considering all the variables except multi-collinear ones, that were excluded from the analysis, based on a correlation coefficient higher than 0.95 (usdm package: Naimi et al., 2014), in order to strengthen the predictor value of the model. Starting from 25 variables, 5 variables were found to have collinearity problem, and were thus excluded from further analyses. The remaining variables were standardised (Hellinger method, Legendre & Gallagher, 2001) and subjected to CCA, with Cultivar, Sex and their interaction as explanatory variables. The CCA result was confirmed through a permutational Multivariate Analysis of Variance (perMANOVA; Anderson, 2001).

The complete R code will be made publicly available on GitHub (https://github.com/matteo-rpm).

## 3. Results

Number of recordings obtained from male and female *S. titanus* adults on the three grapevine cultivars are summarised in Table 2. In particular, 51-cultivar specific recordings were acquired, of which a fraction was selected for further analysis (31 for Barbera, 32 for Brachetto, 37 for Moscato), as described in Material and Methods section.

**Table 2.** Number of total and selected recordings of *S. titanus* feeding behaviour on three grapevine cultivars.

| Cultivar | Total recordings (females, males) | Selected recordings (females, males) |
| --- | --- | --- |
| Barbera | 51 (25, 26) | 31 (18, 13) |
| Brachetto | 51 (25, 26) | 32 (13, 19) |
| Moscato | 51 (25, 26) | 37 (16, 21) |

No significant differences in acquiring successful EPG signals among cultivars, possibly caused by human errors, were found (Pearson’s Chi-squared test, X-squared = 0.36811, df = 2, p-value = 0.8319). From now on, when referring to recordings, only the selected ones will be considered, unless otherwise stated.

Irrespective of the cultivar, most of the insects started probing within the first minute from their access to the leaf (median ± SE = 41 ± 21 s). Behavioral phases were graphically summarised in a temporal progress representation (Figure 1), showing the percentage of leafhoppers in a given phase at different hours. An overall larger area of phloem phase was found for leafhoppers feeding on the Barbera cultivar.

Values of non-phloem variables of all selected recordings are reported in Supplementary File S2. No differences were identified in the variables among the three grapevine cultivars. The proportion of recordings with phloem phases were not significantly different among cultivars (Supplementary File S6, Pearson’s Chi-squared test, X-squared = 0.026378, df = 2, p-value = 0.9869), as almost half of the recordings (45% for Barbera, 44% for Brachetto, 43% for Moscato) showed phloem phases, irrespective of the cultivar (Supplementary File S6). No differences were highlighted among cultivars for all the non-phloem variables (Supplementary File S5), when considering the recordings without a phloem-phase. Further, the non-phloem variables were analysed for recordings with phloem phases (Table 3). Number of events and their duration for the non-phloem phases did not differ among groups (Table 3). Interestingly, the total time spent by the insect with stylets inserted in the plant tissues (“Total probing time”) were similar among the three *Vitis* genotypes. Some differences were found for the related variables “Number of non-probing periods”, and “Number of probes”, as higher values were recorded for both variables on Brachetto, compared to Barbera. On Barbera, females showed fewer “Number of active ingestion (from mesophyll or xylem) phases” than males. No differences were observed between sexes on the other cultivars. No significant differences among cultivars were found for the “Number of phloem ingestions”, or for the “sustained” (longer than 10 minutes) ones (Table 3). Although the “Mean duration of a single event of phloem ingestion” did not differ significantly among cultivars, a longer duration of phloem ingestion events on Barbera was evident. Indeed, significant differences were found for “Total duration of phloem ingestions”, “Duration of the longest phloem ingestion”, and “Time from first probe to first sustained phloem ingestion” between Barbera and the other two grapevine cultivars (Table 3). “Time from first probe to first phloem ingestion” was shorter on Barbera compared to Moscato, with an intermediate duration recorded on Brachetto (Table 3). This also suggests a preference of *S. titanus* for Barbera. For the “Total duration of non-phloematic phases”, for which an effect for the leafhopper sex was found, a difference was recorded between *S. titanus* feeding on Barbera and on Moscato, at least for females. *Scaphoideus titanus* also spent a higher percentage of time in the phloem ingestion phase on Barbera, compared to Brachetto and Moscato cultivars and, consequently, less time in pathway- and active ingestion phases (Table 3). Since the presence of “Np” (typical interruption between two different passive ingestion phases) in phloem phases has been repeatedly recorded (Chuche et al., 2017a; Supplementary File S1 of the present work), three variables were introduced for their description in the present work and are reported in Table 3: “Mean frequency of Np interruptions during phloem phase”, “Percentage of time spent in Np interruption during phloem phase”, and “Number of Np interruptions during phloem ingestion”. The second and third variables showed significant differences between leafhoppers feeding on Barbera and those feeding on the other cultivars, underlying different phloem feeding behaviour on the former cultivar.

**Table 3.**
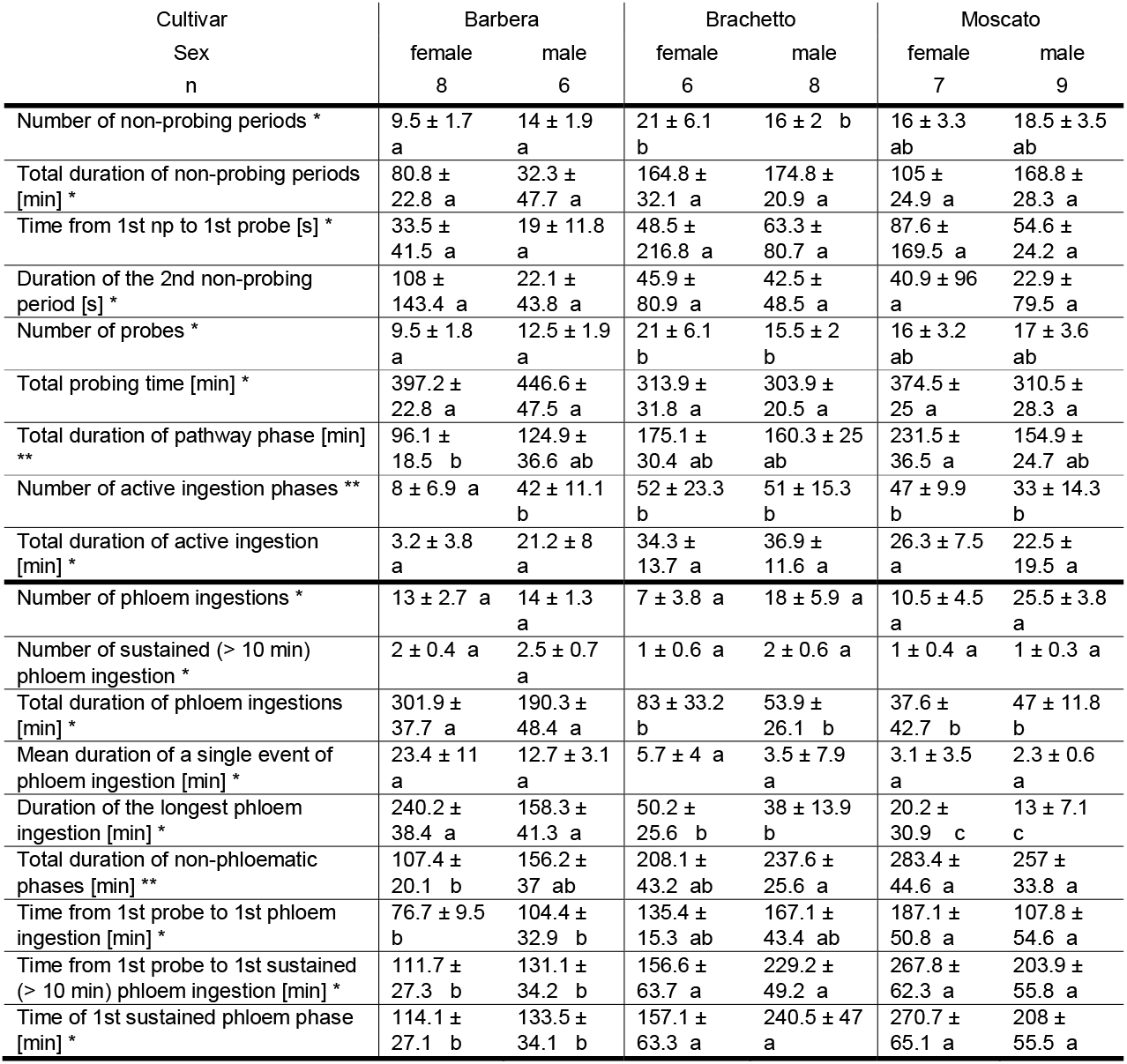
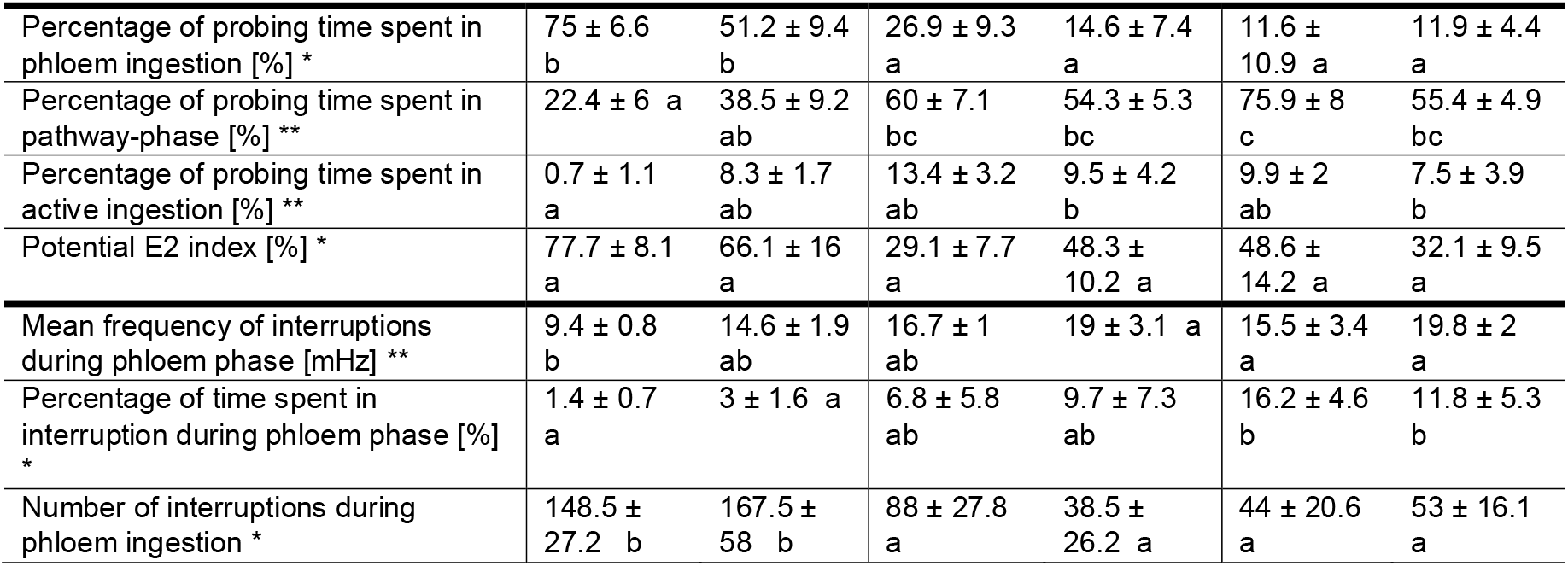
Median ± SE of variables related to recordings presenting phloem phases. Every column reports a single combination of grapevine Cultivar and leafhopper Sex. Every row reports a specific variable. Comparisons between columns were done with a specific GLM family for every variable: quasi-Poisson or negative-binomial for counts, Gamma or inverse-Gaussian for continuous time variables, beta-regression for proportions. Cultivar, Sex and their interaction (Cultivar × Sex) effects for every variable were evaluated. In case of no effect for Sex and Cultivar × Sex, the GLM was run with only Cultivar as explanatory variable (indicated in the tables with the * sign after the specific variable name). In case of effect for Sex or Cultivar × Sex, GLM was run with all the three explanatory variables (indicated in the tables with the ** sign after the specific variable name). Post-hoc comparisons were conducted with least-square means method and Tukey method for p-value adjustment, at significance level as 0.05 and 95% confidence intervals, and represented by letters for every specific group. GLMs specific details (family, coefficients, standard errors, AIC, BIC, R^2^) are reported in Supplementary File S3a-b.

A constrained Canonical Correspondence Analysis (CCA) was conducted to explore the comprehensive effect of the explanatory variables Cultivar, Sex, and their interaction with *S. titanus* feeding behaviour (Figure 2). The CCA is a graphical representation of the non-multi-collinear variables more related to the different groups. In particular, considering the absence of effect for Sex and Cultivar × Sex (Table 4), ellipses were drawn containing 99% confidence intervals for the standard errors related to Cultivar variable. Again, this representation highlighted the difference between *S. titanus* feeding behaviour on Barbera, on one side, and on Brachetto and Moscato, on the other side. Moreover, CCA shows a clear correlation between phloem variables and Barbera cultivar.

**Figure 2.**
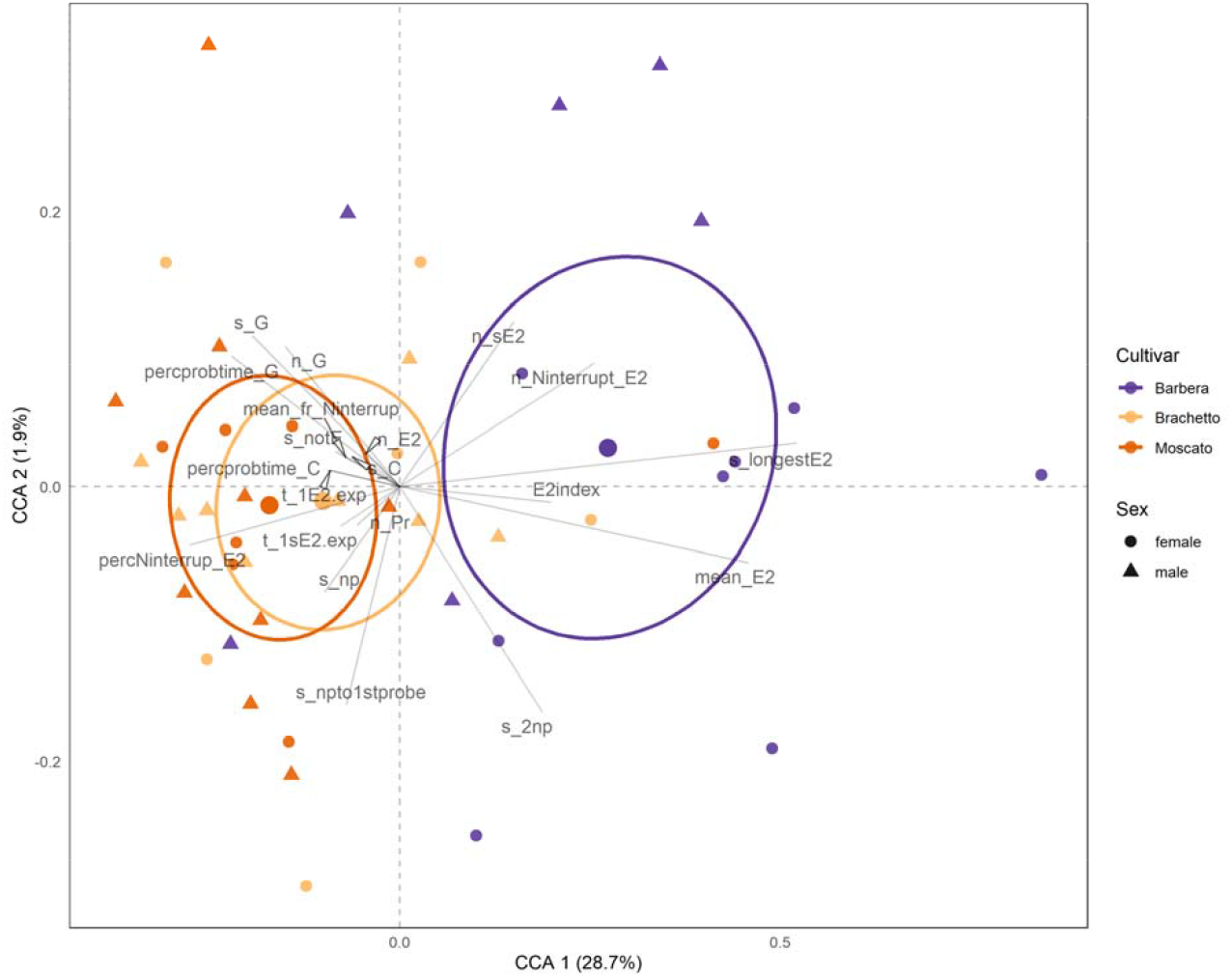
Canonical Correspondence Analysis (CCA) on recordings with phloem phases. The new condensed CCA variables explained 28.7% (CCA 1, x axis) and 1.9% (CCA 2, y axis) of the variability. Cultivar-specific recordings were grouped with ellipses, representing 99% confidence intervals for the standard errors, and the centroid of each was represented. Every point represents a single recording, colour refers to the grapevine cultivar and shape refers to the leafhopper sex. Original variables were plotted and reported with their acronym (Table 1 for acronym interpretation); all variables start from the intersection of the axes and are projected according to their unique composition of CCA 1 and 2.

**Table 4.** perMANOVA results based on Bray-Curtis dissimilarities, using all the non-multi-collinear EPG variables (as described in Materials & Methods section). Df: degrees of freedom; SumOfSqs: sequential sums of squares; F: F statistics values by permutations; Pr(>F): p-values, based on 9999 permutations (the lowest possible p-value is 0.0001).

|  | Df | SumOfSqs | R<sup>2</sup> | F | Pr(>F) | signif |
| --- | --- | --- | --- | --- | --- | --- |
| Cultivar | 2 | 0.21459034 | 0.24924254 | 6.867267 | 0.0001 | *** |
| Sex | 1 | 0.02922068 | 0.03393925 | 1.870225 | 0.1269 |  |
| Cultivar × Sex | 2 | 0.02344144 | 0.02722678 | 0.750167 | 0.6142 |  |
| Residual | 38 | 0.59371753 | 0.68959144 | NA | NA |  |
| Total | 43 | 0.86096998 | 1 | NA | NA |  |

Results of the CCA were confirmed through a perMANOVA (Table 4), which highlighted significant differences among Cultivars, while no significative differences were found for Sex or the interaction of Cultivar and Sex.

## 4. Discussion

In this work, the probing behaviour of the FD leafhopper vector *S. titanus* on grapevine cultivars with different susceptibility to the disease was analysed, to highlight possible differences that can account for different transmission efficiencies. As phytoplasmas are phloem-limited in the plant, vector acquisition and transmission abilities are related to phloem feeding phases, and thus a plant genotype that does not sustain efficient phloem feeding may be less prone to infection.

To understand if probing behaviour of *S. titanus* may contribute to explain tolerance/susceptibility mechanisms of grapevine genotypes, the FD highly susceptible Barbera and the FD tolerant Brachetto and Moscato cultivars (Ripamonti et al., 2021) were compared. Indeed, *S. titanus* showed a feeding preference for the FD highly susceptible Barbera cultivar. To describe *S. titanus*-grapevine interaction, total probing time was subdivided into different behavioral phases, mainly related to the inter/intra-cellular movements of the stylets (pathway-phase), the active ingestion of mesophyll or xylem sap, and the passive ingestion of phloem-sap.

A preference of the leafhopper for Barbera was suggested at first by the overall higher proportion of *S. titanus* feeding on phloem of this cultivar (larger area under the phloem phase, Figure 1), compared to Brachetto and Moscato in the temporal progress area graph. However, it is possible that duration of phloem phases was underestimated in this study, as well as in those of Chuche et al. (2017a, 2017b), and indeed longer recording times can possibly highlight longer durations of phloem ingestion, as hypothesized for *Dalbulus maidis* (Carpane et al., 2011). Under our experimental setting, eight-hour recordings were long enough to allow 50% of the insects to show phloem phase activities, irrespective of the cultivar. This recording time was chosen for the experiments as it represents a widely used standard in EPG studies, and because in previous experiences, Chuche et al. (2017b) showed that, in average, 27% of the *S. titanus* probing time was spent in phloem feeding phases with four hour recordings. According to our results, most of the cultivar-dependent differences in *S. titanus* lies in phloem feeding behaviour. Leafhoppers feeding on Barbera spent more than 50% of their probing time feeding on phloem, while on the other two cultivars the time spent in phloem was less than 20%. This result is in line with an enhanced possibility of acquisition and inoculation of phloem-limited agents, like FDp in the case of Barbera (Galetto et al., 2014). Although the “Potential E2 index”, a parameter regarded as a reliable indicator of phloem acceptability (Alvarez et al., 2006; Girma et al., 1992), was not significantly different among the tested cultivars, higher values were recorded for Barbera, further supporting an improved feeding behavior, that may indicate a better acceptance of the plant as a host. Moreover, since only half of the vectors reached the phloem phase during the 8-hour recordings, we cannot exclude that the amount of time was not sufficient to obtain a more descriptive feeding behaviour from all leafhoppers. Considering the “Time from first probe to first phloem ingestion” some differences were highlighted among cultivars. In particular, a lower amount of time was needed by leafhoppers feeding on Barbera to reach phloem, thus a possible earlier transmission of the phytoplasma to the plant. However, the likelihood of phytoplasma transmission is positively correlated with the length of the inoculation access period (Bosco and Marzachì, 2016), suggesting that the duration of the total phloem activities, rather than its first onset, may be more effective for phytoplasma transmission. Dramatic differences were highlighted in the “Total duration of phloem ingestions” on the different cultivars, while the “Number of phloem ingestions” and “sustained phloem ingestions” were similar. The former was actually the variable accounting for the highest differences in phloem phase among cultivars, and suggests that *S. titanus* prefers Barbera phloem to Brachetto or Moscato ones. Since no differences were recorded among cultivars in the percentage of leafhoppers reaching phloem, but “Total duration of phloem ingestion” and “Duration of the longest phloem ingestion” were higher on Barbera, it can be hypothesized that Brachetto and Moscato phloem saps contain some repellent compounds disturbing phloem feeding. During the phloem phase, the main waveforms were related to the passive ingestion of phloem and to the interruption between two different passive ingestion phases (mainly “Np”). These interruptions were already described for *S. titanus* by Chuche et al. (2017a) and for *Circulifer tenellus* by Stafford and Walker (2009), and were suggested to represent salivation events. Two are the main functions of saliva in piercing-sucking insects: i) production of stylets sheath in the inter-cellular pathway phase (sheath saliva) or ii) dilution of to-be-ingested sap and the suppression of defensive mechanism by the plant through effectors (watery saliva) (Miles, 1972; Tjallingii, 2006; Will, Furch, & Zimmermann, 2013). According to Chuche et al. (2017a) and Stafford & Walker (2009), the “Np” interruptions found during *S. titanus* ingestion of phloem sap correspond to watery-salivation events. This type of salivation is related to the inoculation of persistent-propagative agents from the insect salivary glands into the plants tissues (Hogenhout et al., 2008). Therefore, the greater number of interruption-salivation events on Barbera, that are a reflection of the longer phloem phase, can explain, at least in part, the high susceptibility to FDp of this cultivar. Phytoplasma spread can be regarded as a function of insect acquisition efficiency, which is directly related to the duration of the phloem feeding phase, and of the inoculation efficiency, which is putatively related to the absolute number of watery salivation events, these latter also occuring during phloem feeding phase. According to this hypothesis, on Barbera, the vector acquires and transmits efficiently, because it feeds longer in the phloem and produces a higher number of salivation events compared to Brachetto and Moscato. Indeed, Galetto et al. (2016) demonstrated that FDp acquisition by *S. titanus* depends on the grapevine cultivar, with high efficiency from the most susceptible ones. Also, on Brachetto and Moscato a high frequency of Np interruptions events were recorded, but phloem phase was much shorter, leading to a lower absolute number of salivation events. It is worth noting that, when the three grapevine cultivars were exposed to equally infected leafhoppers, Brachetto and Moscato showed a strong tolerance against the infection (Ripamonti et al., 2021). This is a clear indication that either the inoculation, more than acquisition, has a major impact on transmission efficiency, or plant genotype account for different susceptibilities. The high frequency of Np interruptions on the tolerant cultivars can be explained by the presence of repellent compounds in the phloem saps.

Brachetto and Moscato are aromatic cultivars (Pollon et al., 2019) and they are genetically related (Raimondi et al., 2020). Their leaves contain high quantities of terpenoids (Mazza et al., 2003), and this class of compounds can be transferred through the plant via the phloem flux (Zhang et al., 2016) like other defence compounds (Will et al., 2013). It is possible that *S. titanus* disturbed behavior may be associated with the presence of aromatic compounds, that may act as repellents in Brachetto and Moscato phloems, but further research is needed to clarify if these putative repellent compounds may acts as antixenotic compounds. Antixenosis, defined as the modification of herbivore behaviour by plant factors, which results in the inability of a plant to serve as a host (Kogan and Ortman, 1978; Kordan et al., 2019), is a well-known factor determining host plant resistance. Terpenoids and other volatile and non volatile compounds have well-known antixenotic activities in different plant-insect interactions (Chand et al., 2017; Koul, 2008; Messchendorp et al., 1998). Antixenosis may represent a valuable factor to be considered in the development of grapevine resistance against *S. titanus, de facto* causing a reduction in the inoculation efficiency of FDp. Indeed, leafhopper survival is reduced following a 7 day exposure to Moscato compared to Barbera (Ripamonti et al., 2021). Further research is needed to clarify possible Moscato antibiosis effect on *S. titanus*.

In our study, the leafhoppers started probing within the first minute, regardless of the grapevine cultivar. No evident differences were highlighted in the non-phloem related variables, as well as on total probing time. These results suggest that tested cultivars have no major differences in the biochemical composition or structure of the leaf cuticle, epidermis or mesophyll, that can impact the first feeding behaviour activities. Grapevine trichomes are of the non-glandular type, subdivided in prostrate or erect (Gago et al., 2016). Interestingly, Barbera has a highly dense trichome surface in the abaxial leaf blade (OIV, 2007), suggesting a possible biophysical barrier against piercing-sucking insect nutrition (Smith and Chuang, 2014). Nevertheless, Barbera was the most suitable cultivar for *S. titanus* among those tested. For leafhoppers, data on trichome density acceptability are mainly available for species of the Empoascini tribe, that are mostly insensitive to trichome density on leaves. In fact, *Empoasca vitis* (Pavan and Picotti, 2009), *E. terminalis* (Nasruddin et al., 2014), and *E. fabae* (Kaplan et al., 2009) showed insensitivity to trichome densities on grapevine, soybean, and potato. On the other hand, *E. fabae* and *Amrasca devastans* tend to avoid high trichome density when feeding on edamame (*Glycine max* (L.)) and cotton, respectively (Menger et al., 2018; Murugesan and Kavitha, 2010). Considering that *S. titanus* feeds better on the most highly pubescent cultivar, it can be concluded that a dense abaxially pubescence does not hamper nutrition on grapevine.

This work failed to identify clear differences in feeding behaviour of males and females. Although small differences between sexes were recorded for some variables, no differences were highlighted in the multivariate analysis conducted through CCA followed by perMANOVA. On the contrary, Chuche et al. (2017b) reported that males feed more in the phloem, compared to females. Following the analysis of our EPG recordings, we conclude that no clear differences in feeding behaviour can be identified. Although unlikely, we cannot exclude that the different grapevine cultivars used in the studies may explain for this difference.

Future research should focus on antixenotic compounds in *V. vinifera* genotypes, and their role in vector-associated resistance to FD. On the other hand, plant secondary metabolites involved in defense mechanisms against pathogens, such as polyphenols, particularly vein flavonols and flavanonols (Kedrina-Okutan et al., 2019, 2018) may play a role in plant resistance towards the phytoplasma. All these grapevine genetic traits should be regarded as natural resources to be exploited in the quest for tolerant genotypes for a more sustainable viticulture.

## 5. Conclusions

The feeding behavior profiles of the present work indicate that Barbera cultivar is a better host than Brachetto and Moscato for *S. titanus*. Indeed, the leafhopper showed longer phloem ingestion, with absolute higher number of watery-salivation events, on Barbera grapevines. This feature is consistent with the high susceptibility of Barbera to FDp, as watery salivation has been associated with the inoculation of persistent-propagative agents from the insect salivary glands into the plants tissues. When feeding on Brachetto and Moscato, *S. titanus* showed reduced phloem ingestion of sap, possibly due to antixenotic factors.

## Acknowledgments

This work was funded by the project “Towards resistance to Flavescence dorée of grapevine, Flavoscreen”, Fondazione Compagnia di San Paolo and University of Torino. Authors wish to thank Nicola Bodino for revision of statistical analyses, Marco Chiapello for the fundamental help in the development of the Rwaves package, Stefano Raimondi for suggestions on ampelography, and Giovanni Marchisio for providing grapevine cuttings and dormant wood for *S. titanus* eggs.

## Author contributions

Conceptualization: DB, CM, MR. Data curation: MR. Formal analysis: MR. Funding acquisition: DB, CM. Investigation: MR, FM. Methodology: AF, DC, DB, MR. Software: MR. Project administration: DB. Supervision: AF, DB, DC. Visualization: MR. Writing - original draft: MR. Writing - review & editing: DB, CM, AF, DC.

