## Supplementary File S4 for "*Scaphoideus titanus* Ball feeding behaviour on three grapevine cultivars with different susceptibilities to Flavescence dorée"

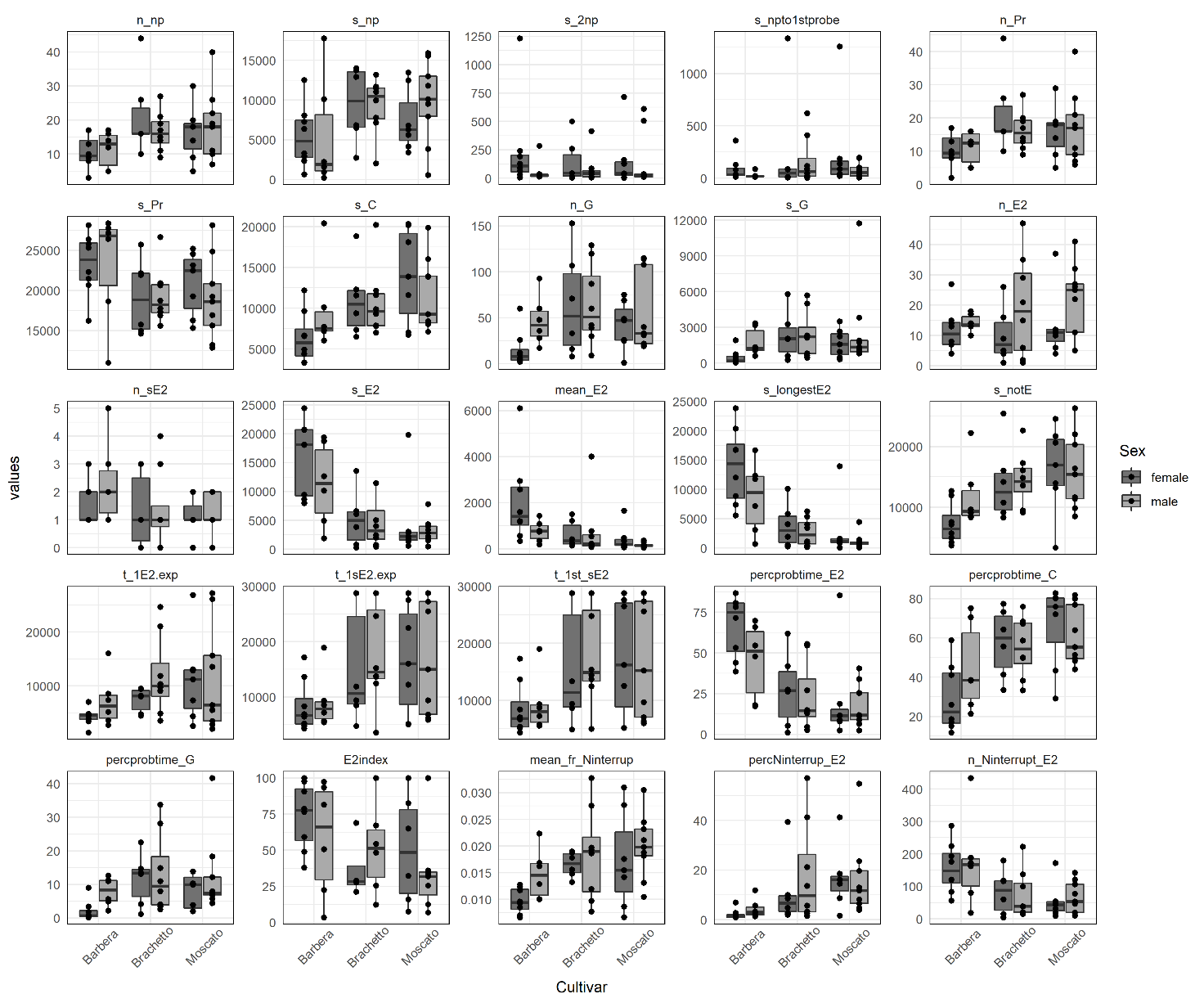

**Supplementary File S4. Graphical representation of EPG variables related to recordings with phloem phases.** For the variable acronyms, see Table 1 in the main text.
