## Supplementary File S5 for "*Scaphoideus titanus* Ball feeding behaviour on three grapevine cultivars with different susceptibilities to Flavescence dorée"

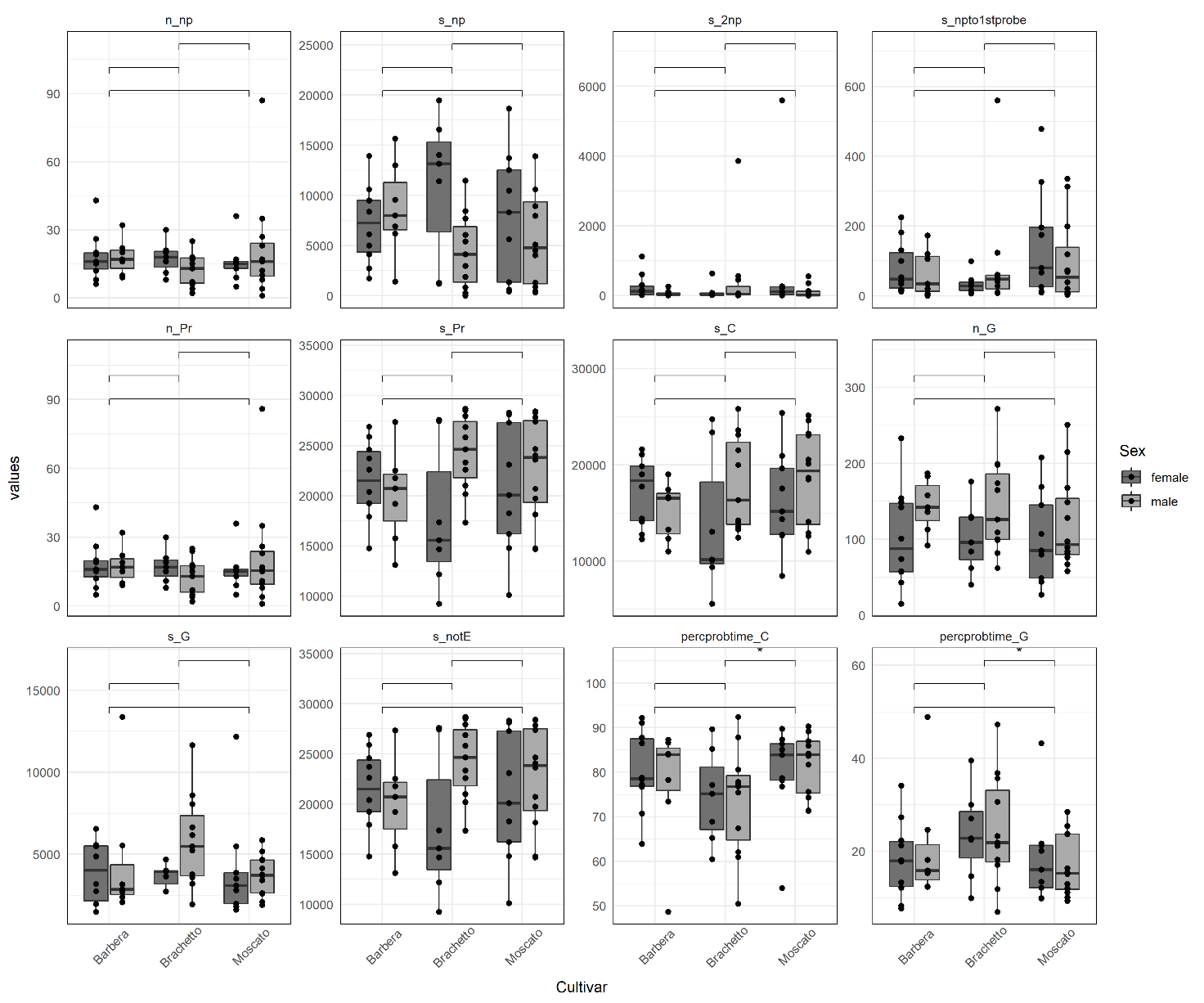

**Supplementary File S5. Graphical representation of EPG variables related to recordings without phloem phases.** Differences between group were evaluated with Wilcoxon rank sum test. For the variable acronyms, see Table 1 in the main text.
