## Supplementary File S6 for "*Scaphoideus titanus* Ball feeding behaviour on three grapevine cultivars with different susceptibilities to Flavescence dorée"

**Supplementary File S6. Number of selected recordings of *S. titanus* probing behaviour with and without phloem phases on three grapevine cultivars.**

| Cultivar | Recordings with phloem (females, males) | Recordings without phloem (females, males) | Percentage of recordings with phloem phase |
| --- | --- | --- | --- |
| Barbera | 14 (8, 6) | 17 (10, 7) | 45.2 |
| Brachetto | 14 (6, 8) | 18 (7, 11) | 43.8 |
| Moscato | 16 (7, 9) | 21 (9, 12) | 43.2 |
