## Supplementary File S1 for "*Scaphoideus titanus* Ball feeding behaviour on three grapevine cultivars with different susceptibilities to Flavescence dorée"

**Supplementary File S1. Summary and graphic representation of the main waveform characteristics for *Scaphoideus titanus* on grapevine.**

| **Waveform** | **Putative plant tissue*** | **Amplitude ± SE [V]** | **Amplitude range (min - max) [V]** | **Amplitude ± SE [%] **** | **Amplitude range (min - max) [%] **** | **Frequency ± SE [Hz]** | **Frequency range (min - max) [Hz]** | **Voltage level ***** |
| --- | --- | --- | --- | --- | --- | --- | --- | --- |
| Pathway-phase | Epidermis and mesophyll | 2.37 ± 0.17 | 0.3 - 5 | 23.68 ± 1.74 | 3 - 50 | Mixed | - | e/i |
| Active ingestion | Mesophyll (< 60 s) and xylem (> 100 s) | 2.41 ± 0.16 | 0.8 - 6.3 | 24.13 ± 1.61 | 8 - 63 | 4.28 ± 0.1 | 2.5 - 6 | e |
| Passive ingestion | Phloem | 0.33 ± 0.03 | 0.07 - 0.9 | 3.31 ± 0.28 | 0.7 - 9 | 3.69 ± 0.13 | 2 - 6 | i |
| "Np" interruption | Phloem | 8.22 ± 0.23 | 4.8 - 11.3 | 82.18 ± 2.28 | 48 - 113 | 25.27 ± 0.34 | 20 - 32 | e/i |

* Putative plant tissue suggested from previous studies (Chuche et al. 2017a; Stafford & Walker, 2009)
** 100% amplitude = 10 V
*** e = extracellular; i = intracellular

Values presented in the table were calculated starting from thirty insect-recordings, ten per cultivar, randomly selected among recordings presenting phloem phases. For each insect, two replicates for the same waveform were randomly selected in all the 8-hour recording, resulting in a total of 60 replicates for every waveform. Examples of the waveform are graphically represented in the figures below.

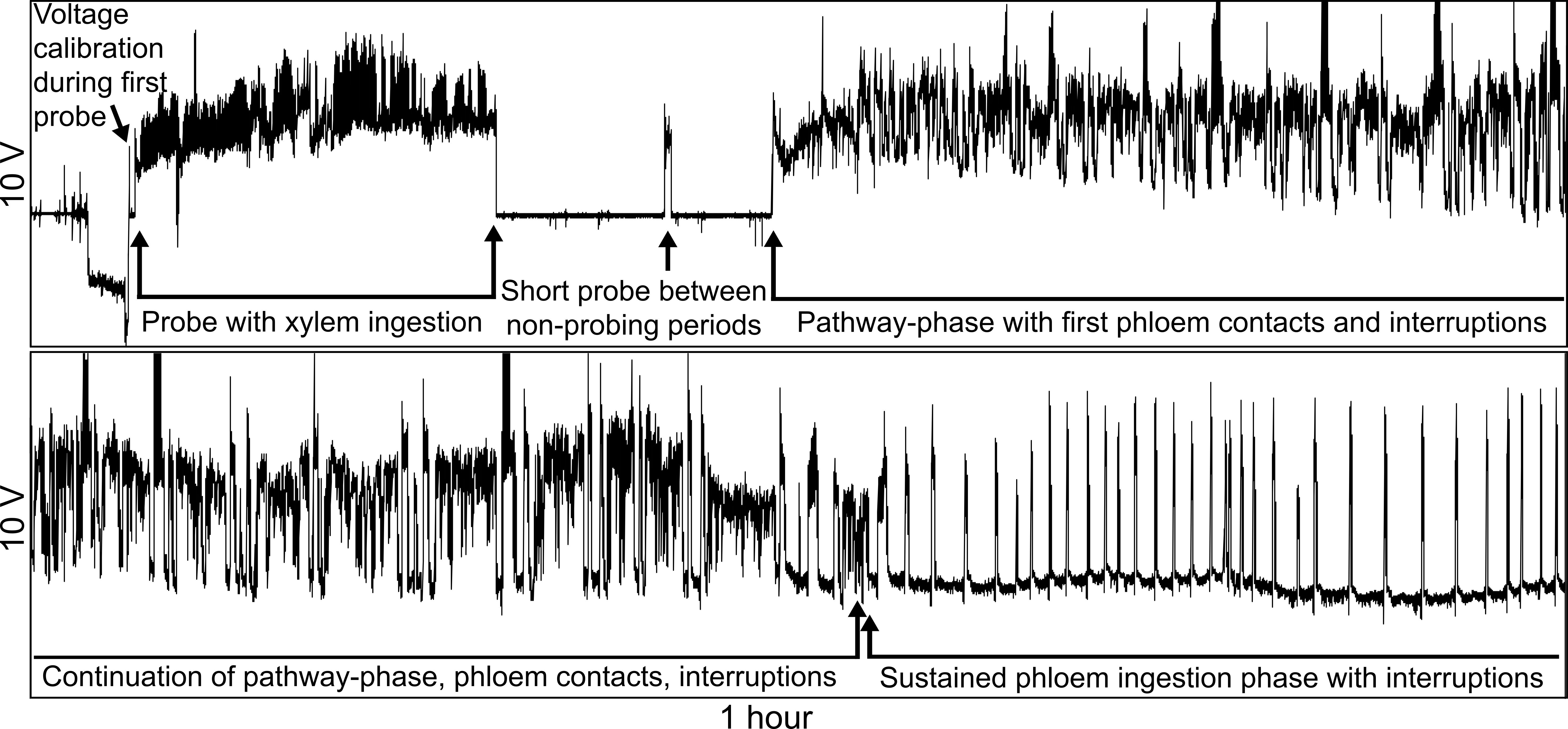

Figure I. Overall view of a 2-hour recording, presenting all the phases described in the table and in the main text.

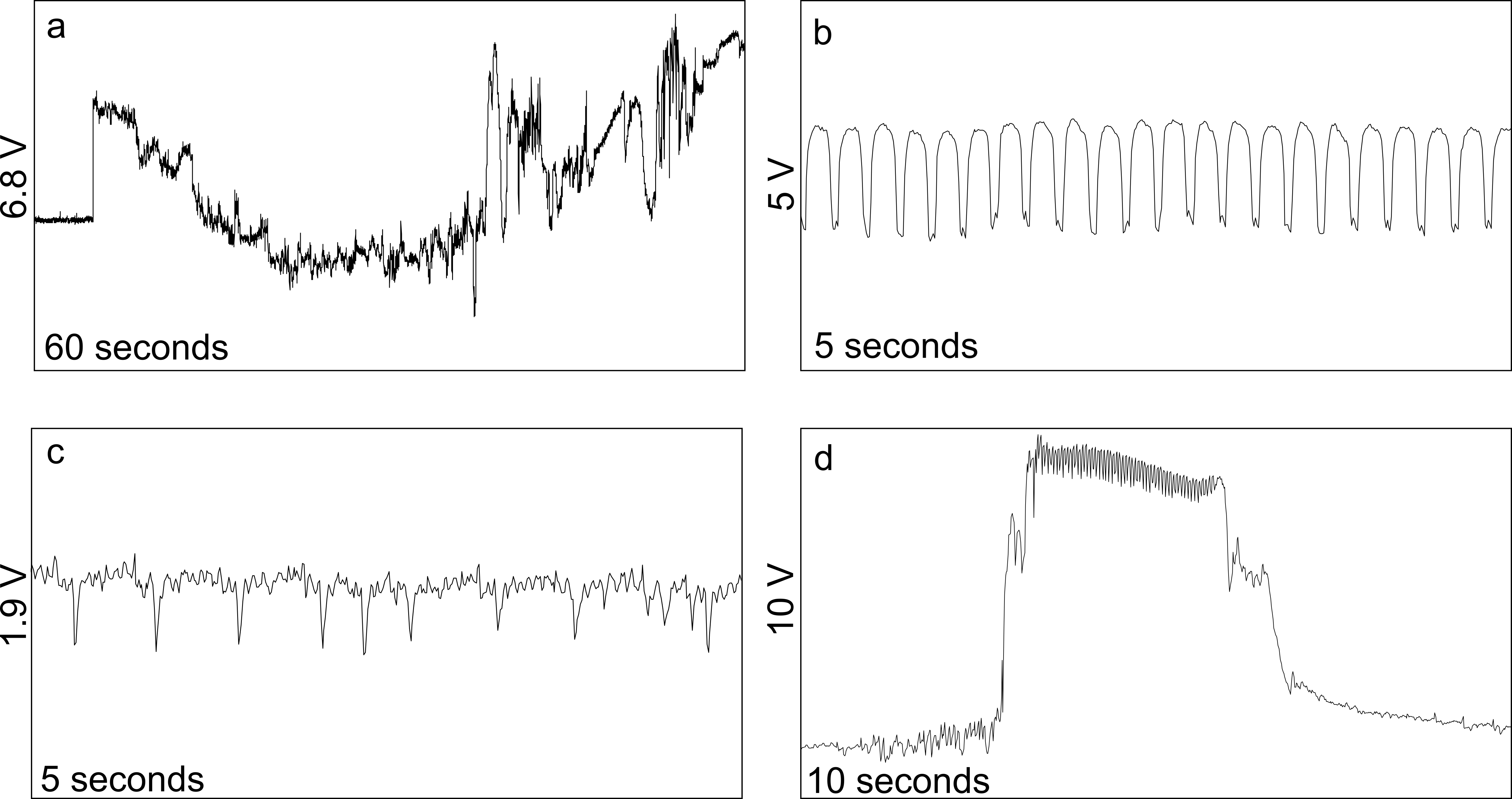

Figure II. Specific waveforms view: a) non-probing, followed by the beginning of a probe as pathway-phase, b) active ingestion of mesophyll or xylem sap, c) passive ingestion of phloem sap, d) “Np” interruption during phloem phases (watery salivation).
