## Supplementary File S2 for "*Scaphoideus titanus* Ball feeding behaviour on three grapevine cultivars with different susceptibilities to Flavescence dorée"

**Supplementary File S2. Median ± SE of non-phloem variables related to all recordings.** Every column reports a single combination of grapevine Cultivar and leafhopper Sex. Every row reports a specific variable. Comparisons between columns were done with a specific GLM family for every variable: quasi-Poisson or negative-binomial for counts, Gamma or inverse-Gaussian for continuous time variables, beta-regression for proportions. Cultivar, Sex and their interaction (Cultivar × Sex) effects for every variable were evaluated. In case of no effect for Sex and Cultivar × Sex, the GLM was run with only Cultivar as explanatory variable (indicated in the tables with the * sign after the specific variable name). In case of effect for Sex or Cultivar × Sex, GLM was run with all the three explanatory variables (indicated in the tables with the ** sign after the specific variable name). Post-hoc comparisons were conducted with least-square means method and Tukey method for p-value adjustment, at significance level as 0.05 and 95% confidence intervals, and represented by letters for every specific group.

| Cultivar | Barbera | Barbera | Brachetto | Brachetto | Moscato | Moscato |
| --- | --- | --- | --- | --- | --- | --- |
| Sex | female | male | female | male | female | male |
| n | 18 | 13 | 13 | 19 | 16 | 21 |
| Number of non-probing periods * | 14 ± 2.4 a | 15 ± 2.1 a | 19 ± 3 a | 14 ± 1.7 a | 16 ± 2.2 a | 17.5 ± 4.2 a |
| Total duration of non-probing periods [min] * | 104.3 ± 15.1 a | 115.3 ± 27 a | 215.2 ± 27.6 a | 119.2 ± 16.5 a | 109.1 ± 22.3 a | 132.8 ± 19.2 a |
| Time from 1st np to 1st probe [s] * | 38.6 ± 22 a | 20.4 ± 15 a | 27.9 ± 100.5 a | 48.4 ± 43.3 a | 83.8 ± 77.8 a | 54.6 ± 21.9 a |
| Duration of the 2nd non-probing period [s] * | 111.7 ± 86.4 a | 24.7 ± 26.3 a | 45.8 ± 57.3 a | 44.3 ± 201.3 a | 100.1 ± 343.9 a | 17.3 ± 43.3 a |
| Number of probes * | 14 ± 2.4 a | 15 ± 2 a | 17 ± 2.9 a | 14.5 ± 1.8 a | 16 ± 2.2 a | 15.5 ± 4.3 a |
| Total probing time [min] * | 374.7 ± 15.1 a | 363.2 ± 26.9 a | 262.5 ± 27.5 a | 350.2 ± 16.6 a | 354.8 ± 22.3 a | 345.5 ± 19.3 a |
| Total duration of pathway phase [min] * | 209 ± 25.4 a | 205.9 ± 22.8 a | 169.7 ± 28.3 a | 226.8 ± 22.3 a | 246.4 ± 22.4 a | 235.2 ± 21.3 a |
| Number of active ingestion phases * | 50 ± 15.8 a | 93 ± 16.3 a | 84 ± 14.7 a | 100 ± 15.5 a | 69 ± 14.6 a | 84 ± 13.8 a |
| Total duration of active ingestion [min] * | 32.5 ± 8.9 a | 45 ± 15.2 a | 54.4 ± 7.2 a | 63.1 ± 11.4 a | 40.9 ± 11.6 a | 44 ± 8.9 a |
